# Dalpiciclib and Pyrotinib Exert Synergistic Antitumor Effects in Triple Positive Breast Cancer

**DOI:** 10.1101/2021.10.04.463019

**Authors:** Jiawen Bu, Yixiao Zhang, Nan Niu, Kewei Bi, Lisha Sun, Xinbo Qiao, Yimin Wang, Yinan Zhang, Xiaofan Jiang, Dan Wang, Qingtian Ma, Huajun Li, Caigang Liu

## Abstract

**Background:** The therapeutic benefit of the standard combination of anti-HER2 and chemotherapy in triple-positive breast cancer (TPBC) is limited even after the addition of endocrine therapy to the regimen. Therefore, treatment optimization is required urgently.

**Methods:** Through the drug sensitivity test, the drug combination efficacy of anti-HER2 drug, endocrine drug and CDK4/6 inhibitor to BT474 cells were tested. The underlying molecular mechanisms were investigated using immunofluorescence, western blot analysis, immunohistochemical staining and cell cycle analysis. Potential biomarker which may indicate the responsiveness to drug treatment in triple positive breast cancer was selected out using RNA-sequence and tested using immunohistochemical staining.

**Results:** We found that pyrotinib combined with dalpiciclib showed better efficacy than pyrotinib combined with tamoxifen in BT474 cells. Degradation of HER2 could enhance ER nuclear transportation, whereas cell cycle blockers could reverse this process. This may be the underlying mechanism by which the addition of dalpiciclib was more beneficial than the addition of pyrotinib plus tamoxifen. Furthermore, CALML5 was revealed to be a potential indicator of responsiveness to anti-HER2 therapy plus CDK4/6 inhibition in triple positive breast cancer.

**Conclusion:** Our study provided evidence for the introduction of CDK4/6 inhibitor in the treatment of TPBC and indicated that the combination of anti-HER2 therapy and cell cycle blockers may be a better strategy for TPBC treatment.

**Funding:** This study was supported by the National Natural Science Foundation of China (#U20A20381, #81872159)

## Introduction

Human epidermal growth factor receptor 2-positive (HER2^+^) breast cancer is associated with an increased risk of disease recurrence and death (*Perou et al., 2000; Slamon et al., 1987; Tzahar et al., 1996*). HER2-overexpressing breast tumors have high heterogeneity, accounting partially for the co-expression of hormone receptors (HR) (*Loi et al., 2016*). Previous studies have demonstrated that extensive cross-talk exists between the HER2 signaling pathway and the estrogen receptor (ER) pathway (*Wang et al., 2011*). In addition, exposure to anti-HER2 therapy may reactivate the ER signaling pathway, which could lead to drug resistance (*Brandao et al., 2020*). Generally, however, HER2-positive patients are treated using the same algorithms, both in the early and advanced stages (*Moja et al., 2012*) Thus, novel therapeutic strategies are urgently needed for patients with HER2^+^/HR^+^ breast cancer.

Increasing evidence has confirmed that the intrinsic differences between HER2^+^/HR^+^ and HER2^+^/HR^-^ patients should not be ignored (*Carey et al., 2016*). Clinical outcomes have demonstrated that HER2^+^/HR^+^ breast cancer patients have a lower chance of achieving a pathologically complete response than HER2^+^/HR^-^patients, when treated with neoadjuvant chemotherapy plus anti-HER2 therapy (*Cameron et al., 2017; Cortazar et al., 2014*). Nevertheless, the addition of concomitant endocrine therapy to anti-HER2 therapy or chemotherapy did not show any advantages in clinical trials, such as the NSABP B-52 and ADAPT HER2^+^/HR^+^studies (*Harbeck et al., 2017; Rimawi et al., 2017*). Therefore, whether endocrine therapy is useful in HER2^+^/HR^+^ breast cancer treatment is questionable.

Recently, the synergistic effect of CDK4/6 (cyclin kinase 4/6) inhibitors and anti-HER2 drugs in HER2^+^ breast cancer has been reported. The combination of anti-HER2 drugs and CDK4/6 inhibitors showed strong synergistic effects and high efficacy in HER2^+^ breast cancer cells (*Goel et al., 2016; Zhang et al., 2019*). The combination of CDK4/6 inhibitors and HER2-targeted therapy as an alternative strategy for HER2^+^/HR^+^ patients, warrants further exploration.

Herein, we investigated the combined effect of pyrotinib (anti-HER2 drug), tamoxifen (endocrine therapy), and dalpiciclib (CDK4/6 inhibitor) on the triple-positive breast cancer (TPBC) cell line BT474. We found that pyrotinib combined with dalpiciclib showed better efficacy than pyrotinib combined with tamoxifen. In addition, HER2-targeted therapy induced nuclear ER redistribution in TPBC cells, which could be reversed by the addition of a CDK4/6 inhibitor. Furthermore, CALML5 could be a potential indicator of responsiveness to HER2-targeted therapy combined with a CDK4/6 inhibitor. Our study provided a new function of the CDK4/6 inhibitor in TPBC cells treated with anti-HER2 therapy and suggested a novel strategy for improving the clinical response in TPBC.

## Results

### Pyrotinib combined with dalpiciclib shows better efficacy than when combined with tamoxifen

To explore the effects of pyrotinib, tamoxifen, and dalpiciclib in TPBC, we first evaluated the cytotoxic activities of these three reagents in BT474 breast cancer cells. The results showed that the IC50 doses for pyrotinib, tamoxifen, and dalpiciclib were 10 nM, 5 μM, and 8 μM, respectively (Figure 1-figure supplement 1a). To further investigate whether these three drugs could have a synergistic effect in BT474 cells, we assessed the efficacies of the combinations of pyrotinib and dalpiciclib, pyrotinib and tamoxifen, and tamoxifen and dalpiciclib on the inhibition of cell proliferation at different concentrations. We calculated the combination index for each combination using Compusyn software to determine if the antitumor effects were synergistic (*Chou and Talalay, 1984*). Synergistic effects were observed in the combination group of pyrotinib and dalpiciclib, as well as in the pyrotinib and tamoxifen groups; both with CI values of <1 at several concentrations (Figure 1a). However, in the combination group of tamoxifen and dalpiciclib, no synergistic effect was observed.

**Figure 1.**
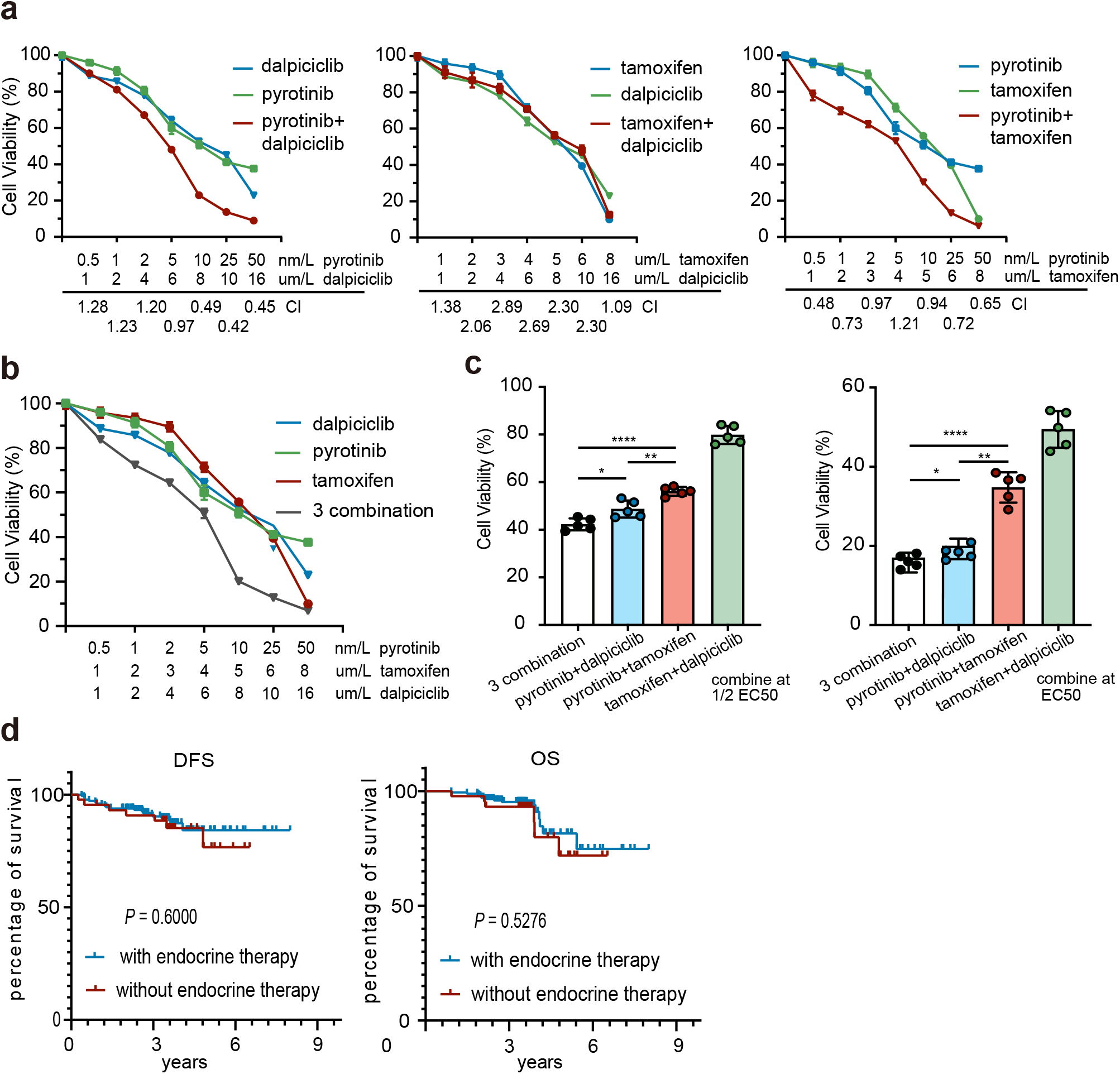
Drug sensitivity test and cell cycle analysis of pyrotinib, tamoxifen, dalpiciclib and their combination on BT474 cells. a-b: Drug sensitivity assay of BT474 cells to single drug and different drug combination (Data was presented as mean ± SEMs, all drug sensitivity assay were performed independently in triplicates). c: Drug sensitivity assay of BT474 cells to different drug combination at EC50 concentration and 1/2 EC50 concentration. (Data was presented as mean ± SDs, \**P*<0.05, \*\**P*<0.01 and \*\*\*\**P*<0.0001 using Student’s *t*-test; all the assays were performed independently in triplicates). d: DFS (*P*=0.600) and OS (*P*=0.5276) analysis of HER2+/HR+ patients received adjuvant therapy with (n=177) or without endocrine therapy (n=44). *P*-values were obtained by log-rank test.

We also analyzed the effect of the three-drug combination, and it showed a stronger cytotoxic effect on TPBC compared with the effect of the other two-drug combinations (Figure 1b). As both dalpiciclib and tamoxifen showed synergistic effects in combination with pyrotinib, we sought the combination that showed better efficacy. Hence, we treated the BT474 cells with different combinations at EC50 or half EC50 concentrations. The three-drug combination and the combination of pyrotinib and dalpiciclib showed a stronger cell inhibition compared with that exerted by pyrotinib and tamoxifen as well as tamoxifen and dalpiciclib (Figure 1c). The colony formation assay also showed similar trends as the cell viability assay; the three-drug combination formed the least number of colonies, followed by the combination of pyrotinib and dalpiciclib (Figure 1-figure supplement 1b-c).

To verify the efficacy of endocrine therapy in HR^+^/HER2^+^ patients, we retrospectively analyzed the clinical data of 221 HR^+^/HER2^+^ patients who received adjuvant therapy at the Shengjing Hospital. Of these, 44 patients received anti-HER2 therapy plus chemotherapy, and 177 patients received anti-HER2 therapy combined with chemotherapy and endocrine therapy. A Kaplan-Meier analysis showed that the addition of endocrine therapy to adjuvant anti-HER2 therapy plus chemotherapy did not significantly alter disease-free survival (DFS; *P* = 0.600) or overall survival (OS; *P* = 0.5276) in HR^+^/HER2^+^ patients (Figure 1d).

### Nuclear ER distribution is increased after Anti-HER2 therapy and could be reversed by the introduction of a CDK4/6 inhibitor

The results of the drug sensitivity test showed that the combination of pyrotinib and tamoxifen was less effective than the combination of pyrotinib and dalpiciclib. Considering that anti-HER2 therapy may activate the ER signaling pathway, we performed immunofluorescence staining for ER distribution on the different drug-treated groups. We found that pyrotinib induced ER nuclear translocation in BT474 cells, which could be partially reversed by the addition of dalpiciclib, rather than tamoxifen (Figure 2a). To further investigate whether the expression of HER2 could affect the distribution of ER, we transfected MCF7 cells with HER2 overexpression plasmids. We found that ER in MCF7 cells (WT and NC) was mainly expressed in the nuclei, whereas in the HER2 overexpressing MCF7 cells, ER was distributed throughout the cell plasma. Treating HER2 overexpression MCF7 cells with pyrotinib could redistribute the ER into the nuclei (Figure 2b). Western blot analyses revealed although the nuclear ER levels increased considerably, the total expression of ER was reduced after the use of pyrotinib (Figure2-figure supplement 1a-b). The use of tamoxifen increased the expression of total ER and nuclear ER, significantly, while the expression of nuclear ER in combination with pyrotinib and dalpiciclib showed a decrease (Figure2-figure supplement 1a-b).

**Figure 2.**
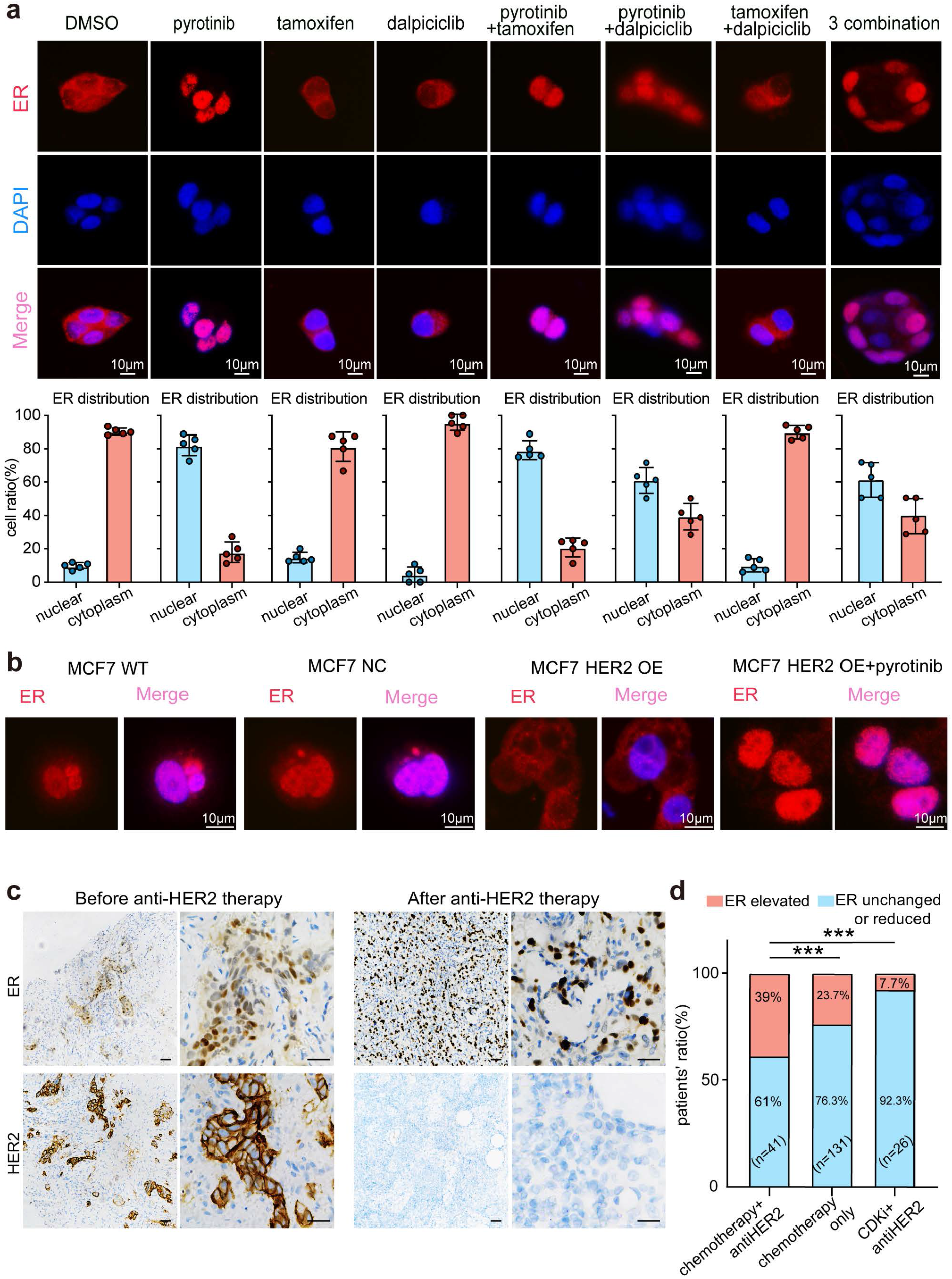
Anti-HER2 therapy could lead ER shifting into cell nucleus in HER2^+^/HR^+^ breast cancer while CDK4/6 inhibitor could reverse the nuclear translocation of ER. a: Distribution of estrogen receptor in BT474 cell line after different drug treatment. (The distribution ratio of ER was calculated manually by randomly chosen 5 views in 400magnification. All the assays were performed independently in triplicates). b: Distribution of estrogen receptor in wild type MCF7 cells, MCF7 cells transfected with HER2 over-expression plasmids and MCF7 cells transfected with HER2 over-expression plasmids treated with pyrotinib. (All the assays were performed independently in triplicates). c: Representative views of ER and HER2 expression in patients before and after anti-HER2 neoadjuvant therapy. d: Ratio of patients with elevated ER expression and patients with unchanged or reduced ER expression in different kinds of neoadjuvant therapy groups. (\*\*\**P*<0.001 using Student’s *t*-test)

Based on our *in vitro* findings, we further explored the ER distribution in clinical samples from the different treatment groups. To this end, we collected the clinical information of 172 HER^+^/HR^+^ patients who received neoadjuvant therapy at the Shengjing Hospital (Table 1). We found significant elevations in the nuclear ER expression levels of patients who received chemotherapy and anti-HER2 therapy, compared with the levels in patients who only received chemotherapy (Figure 2c, d). However, in our ongoing clinical trial (NCT04486911), the nuclear ER expression levels of patients did not show significant elevations after the HER2-targeted therapy combined with dalpiciclib (Figure 2d). These findings verified that the ER receptor may have shifted to the nucleus after anti-HER2 therapy, which could be reversed with the introduction of a CDK4/6 inhibitor.

**Table 1.** Demographic information of triple positive breast cancer patients who received neoadjuvant therapy.

| Variables | Chemotherapy | Chemotherapy+antiHER<br>2 therapy | *p*-value |
| --- | --- | --- | --- |
| No.of patients | 131 | 41 |  |
| Age of year |  |  | ns |
| ≤50 | 82(62.60) | 25(61.00) |  |
| >50 | 49(37.40) | 16(39.00) |  |
| T stage |  |  | ns |
| 1 | 15(11.45) | 5(12.20) |  |
| 2 | 90(68.70) | 32(78.04) |  |
| 3 | 26(19.85) | 4(9.76) |  |
| ER status |  |  | ns |
| ≤30% | 31(23.66) | 8(19.51) |  |
| >30% | 100(76.34) | 33(80.49) |  |
| PR status |  |  | 0.0059 |
| ≤30% | 80(61.07) | 15(36.59) |  |
| >30% | 51(38.93) | 26(63.41) |  |
| HER2 status |  |  | 0.0007 |
| (++) | 78(59.54) | 12(29.27) |  |
| (+++) | 53(40.46) | 29(70.73) |  |
| Ki67 index |  |  | ns |
| <20% | 51(38.93) | 16(39.00) |  |
| >20% | 80(61.07) | 25(61.00) |  |

### Bioinformatic analyses unravel the synergistic mechanisms underlying the dalpiciclib and Anti-HER2 therapy in TPBC

To further explore the synergistic mechanisms of the addition of CDK4/6 inhibitor treatment in HER2^+^/HR^+^ breast cancer, we first analyzed the gene expression profiles of the breast tumor cells treated with pyrotinib via RNA-seq. The signaling pathway enrichment analysis of the differentially expressed genes (DEGs) showed that majority of the DEGs were significantly enriched in the TNF signaling pathway and cell cycle, while steroid biosynthesis was also strongly active, suggesting that the steroid hormone pathway was activated by pyrotinib (Figure 3a-b). Similar results were obtained from the Gene Set Enrichment Analysis (GSEA). The administration of pyrotinib resulted in downregulation of the cell cycle and activation of the hormone pathway. The leading-edge subset of these pathways included the MITOTIC SPINDLE, G2M CHECKPOINT, and ESTROGEN RESPONSE EARLY (Figure 3c). These results showed good concordance with our *in vitro* findings.

**Figure 3.**
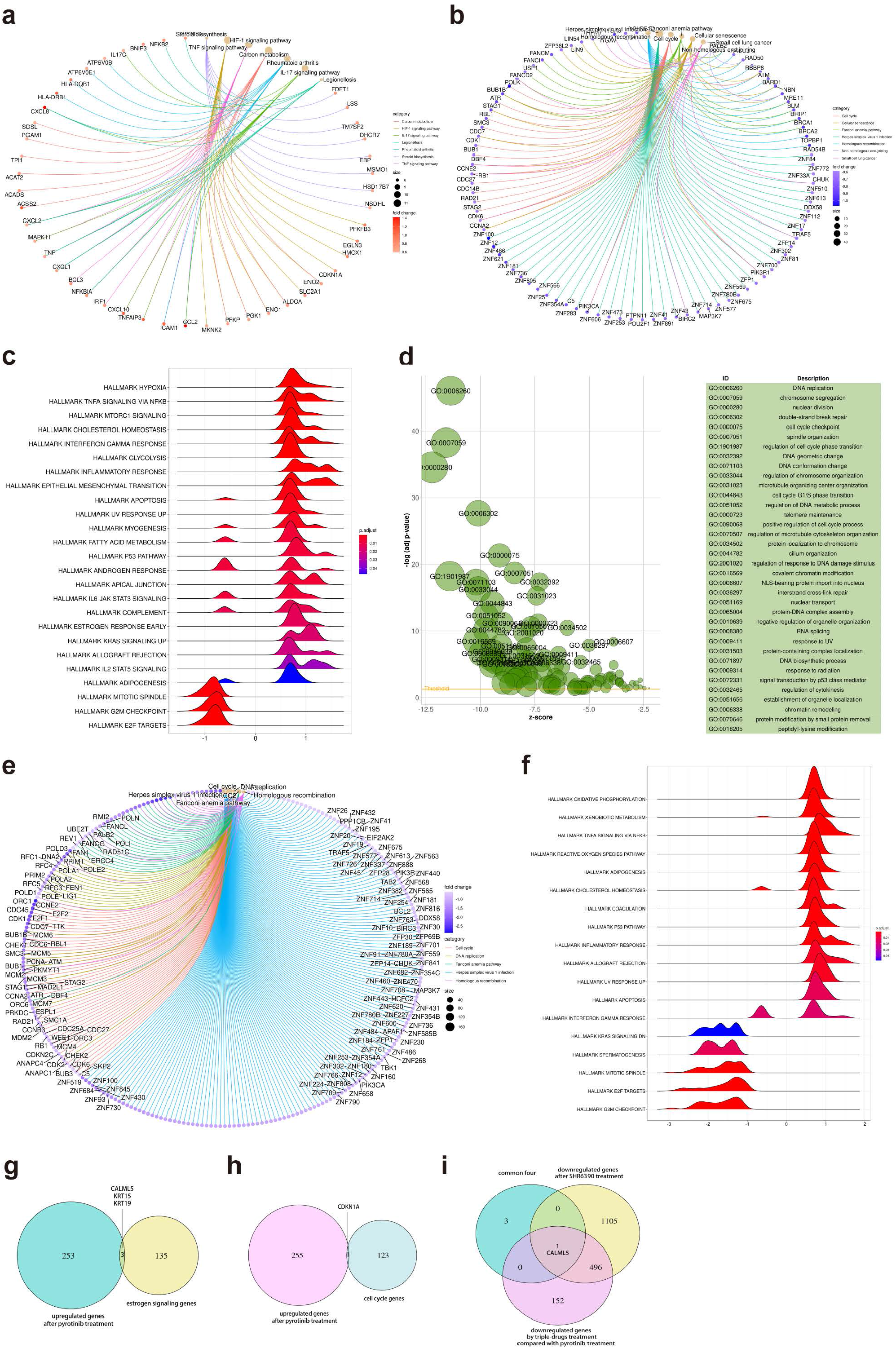
Bioinformatic analysis revealed dalpiciclib and pyrotinib blocking HER2 pathway and cell cycle in BT474 cells synergistically. a-b: Signaling pathway enrichment analysis of mRNA changes of BT474 cells treated with pyrotinib compared to BT474 cells treated 0.1%DMSO c: GSEA analysis of mRNA changes of BT474 cells treated with pyrotinib compared to BT474 cells treated 0.1%DMSO. d-e: Signaling pathway enrichment analysis of mRNA changes of BT474 cells treated with pyrotinib+ tamoxifen+dalpiciclib compared to BT474 cells treated with pyrotinib+ tamoxifen. f: GSEA analysis of mRNA changes of BT474 cells treated with pyrotinib+ tamoxifen+ dalpiciclib compared to BT474 cells treated with pyrotinib+ tamoxifen. g: Intersection of genes which was upregulated after pyrotinib treatment and belonged to estrogen receptor signaling pathway. h: Intersection of genes which was upregulated after pyrotinib treatment and belonged to cell cycle genes. i: Intersection of genes which was upregulated after pyrotinib treatment and was downregulated after the introduction of dalpiciclib.

We then investigated the alteration of the gene expression profiles between breast tumor cells treated with triple-combined drugs (pyrotinib, tamoxifen, and dalpiciclib) and those treated with the dual-combined drugs (pyrotinib and tamoxifen) via gene enrichment analyses. The results suggested that the addition of dalpiciclib markedly reduced cell cycle progression. This was characterized by the enrichment of the cell cycle and the DNA replication process (Figure 3e). The GSEA results further indicated that the progression of the cell cycle was impeded by the enrichment of the gene sets, including MITOTIC SPINDLE and G2M CHECKPOINT (Figure 3f).

The inhibition of the ER pathway might be involved in the effect of pyrotinib plus dalpiciclib on TPBC cells; therefore, intersection analyses were performed to confirm this. As shown in Figure 3g, *CALML5*, *KRT15*, and *KRT19* are the common genes shared between the two sets, the upregulated genes treated with pyrotinib and the genes belonging to the estrogen signaling pathway. Since dalpiciclib is a cell cycle blocker, we also analyzed the common genes involved in the upregulation of the genes and the cell cycle progression after pyrotinib treatment. *CDKN1A* was the only shared gene in these two sets (Figure 3h). We then investigated whether any of the above-mentioned genes were upregulated with the use of pyrotinib and whether this could be reversed with the introduction of dalpiciclib, which may serve as a potential biomarker of the responsiveness to different treatments. The results showed that only one factor, *CALML5,* was the common gene (Figure 3i).

### CALML5 is a potential indicator for the responsiveness to anti-HER2 therapy plus CDK4/6 inhibitor

Western blot analyses and bioinformatic analyses were conducted to verify the changes in the signaling pathways. The western blot analyses showed that while the introduction of tamoxifen did not significantly affect the expression of HER2 and partially inhibited the HER2 downstream pathway (AKT-mTOR signaling pathway), it did not significantly affect the phosphorylation of Rb (Figure 4a). In contrast, after the introduction of dalpiciclib, the activation of mTOR was partially inhibited, which relieved the negative feedback on the HER2 pathway, as evidenced by the slight increase in the HER2 and pAKT, which maintained the sensitivity of the HER2 pathway to pyrotinib. This was consistent with the findings of Goel et al (*Goel et al., 2016*). The combination of pyrotinib and dalpiciclib significantly reduced pRb expression (Figure 4a). In addition, cell arrest analyses of the different drug combinations were performed. As shown in Figure 4b, compared with the cells treated with pyrotinib or tamoxifen, the introduction of dalpiciclib significantly increased the number of cells arrested in the G1/S phase. This confirmed the synergistic inhibition of cell proliferation by dalpiciclib and pyrotinib.

**Figure 4.**
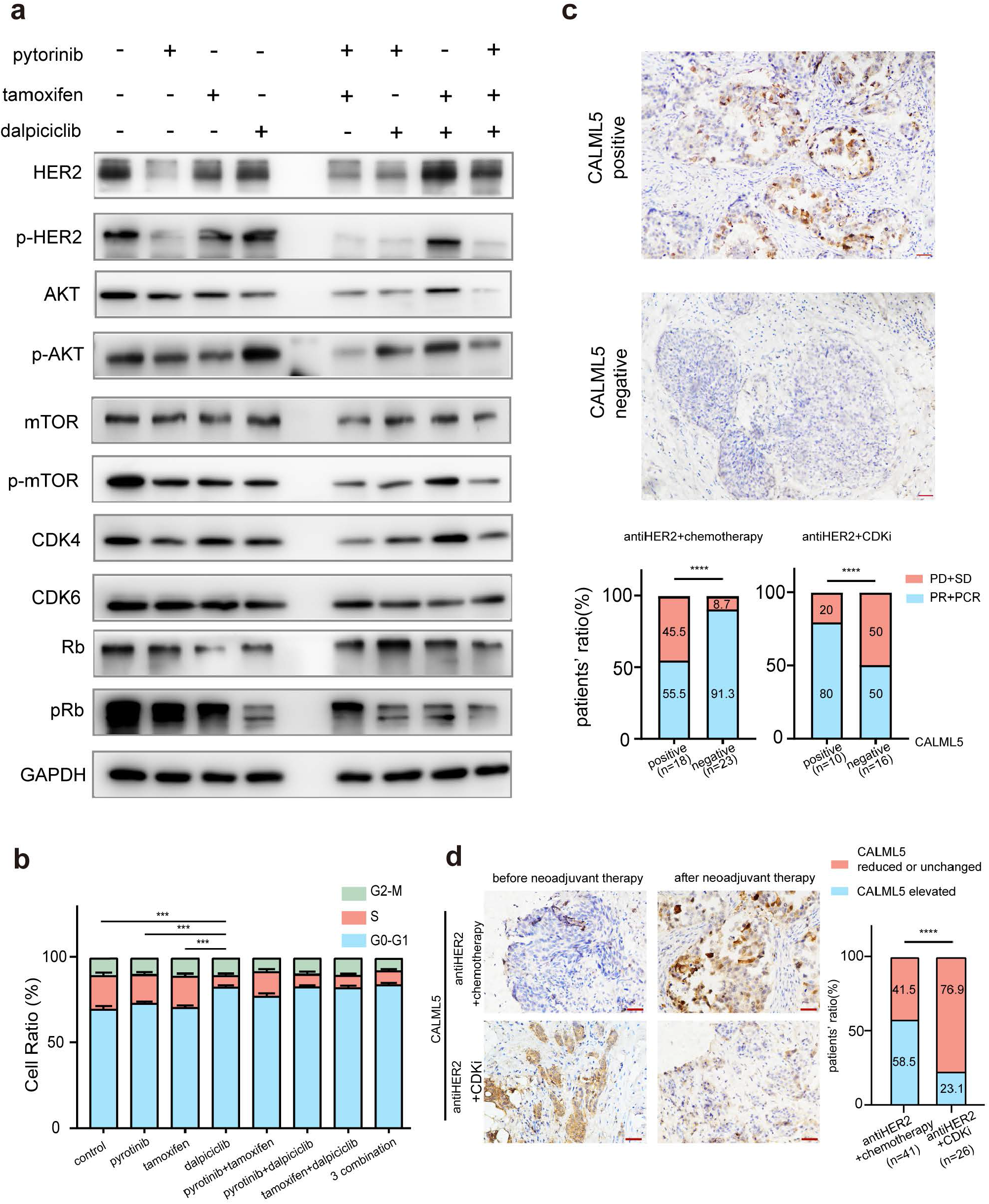
CALML5 could serve as a predictor for responsiveness of anti-HER2 therapy plus CDK4/6 inhibitor. a: Western blot analysis of HER2 signaling pathway and cell cycle pathway in BT474 cells treated with different drugs or their combination. (This assay was performed in triplicates independently). b: Cell cycle analysis in BT474 cells treated with different drugs or their combination. (Data was presented as mean ±SDs, \*\*\**P*<0.001 using Student’s *t*-test; all the assays were performed independently in triplicates). c: Representative views of CALML5 positive/negative tissue. The difference of PR+PCR ratio and PD+SD ratio in patients who received anti-HER2 therapy plus chemotherapy or anti-HER2 therapy plus CDK4/6 inhibitor regarding on their expression of CALML5. (\*\*\*\**P*<0.0001 using Student’s *t*-test). d: Representative views of CALML5 positive/negative tissue. Ratio of patients with elevated or decreased CALML5 after receiving anti-HER2 therapy plus chemotherapy, or anti-HER2 therapy plus CDK4/6 inhibitor. (\*\*\*\**P*<0.0001 using Student’s *t*-test).

To verify whether CALML5 could be a potential predictor of treatment responsiveness in clinical practice, clinical biopsy samples were collected from HER2^+^/HR^+^ patients before and after neoadjuvant therapy (anti-HER2 therapy plus chemotherapy or anti-HER2 therapy plus CDK4/6 inhibitor). The demographic information is shown in Table 2. Immunohistochemical staining of CALML5 showed that the CALML5-positive cells indicated better drug sensitivities and higher probabilities of achieving pathological complete response (pCR) and PR in patients receiving anti-HER2 therapy and a CDK4/6 inhibitor compared with anti-HER2 therapy plus chemotherapy (Figure 4c). Moreover, the positive rate of CALML5 decreased after anti-HER2 therapy plus a CDK4/6 inhibitor treatment (Figure 4d), consistent with the results of the bioinformatic analyses.

**Table 2.** Demographic information of triple positive breast cancer patients who were tested for CALML5 before receiving neoadjuvant therapy.

| Variables | Chemotherapy+anti<br>HER2 therapy | Pyrotinib+dalpiciclib+ta<br>moxifen | *p*-value |
| --- | --- | --- | --- |
| No.of patients | 41 | 26 |  |
| Age of year |  |  |  |
| ≤50 | 25(61.00) | 16(61.53) | ns |
| >50 | 16(39.00) | 10(38.47) |  |
| T stage |  |  |  |
| 1 | 5(12.20) | 2(7.70) | ns |
| 2 | 32(78.04) | 21(80.76) |  |
| 3 | 4(9.76) | 3(11.54) |  |
| ER status |  |  |  |
| ≤30% | 8(19.51) | 2(7.6) | 0.0145 |
| >30% | 33(80.49) | 24(92.4) |  |
| PR status |  |  |  |
| ≤30% | 15(36.59) | 13(50) | ns |
| >30% | 26(63.41) | 13(50) |  |
| HER2 status |  |  |  |
| (++) | 12(29.27) | 10(38.5) | ns |
| (+++) | 29(70.73) | 16(61.5) |  |
| Ki67 index |  |  |  |
| <20% | 16(39.00) | 8(30.8) | ns |
| >20% | 25(61.00) | 18(69.2) |  |
| CALML5 |  |  |  |
| positive | 31(75.61) | 19(73.08) | ns |
| negative | 10(24.39) | 7(26.92) |  |

ns, not significant.

## Discussion

Until now, the combination of antiHER2 therapy and chemotherapy have been the major treatment strategies for treatment of TPBC, although sometimes this is combined with endocrine therapy (*Gianni et al., 2012; Schneeweiss et al., 2013*). Although pCR and DFS improve with the use of the combination of anti-HER2 therapy and chemotherapy, the strong adverse effects of chemotherapy cannot be ignored (*Maguire et al., 2021*). Moreover, clinical data showed that the addition of anti-estrogen receptor drugs in the treatment regimen of TPBC did not provide additional advantages in the pCR rates and DFS (*Harbeck et al., 2017; Rimawi et al., 2017*). Hence, with the rapid development of small-molecule drugs such as tyrosine kinase inhibitors (TKIs) and CDK4/6 inhibitors, additional chemo-free strategies are being developed for the treatment of HER2^+^/HR^+^ breast cancer (*Gianni et al., 2018; Pascual et al., 2021; Saura et al., 2014*). In the present study, we investigated whether the addition of CDK4/6 inhibitors to HER2-targeted and endocrine therapy could be an alternative strategy for HER2^+^/HR^+^ patients.

In our study, we found that the combination of tamoxifen and pyrotinib was less effective than the combination of pyrotinib and dalpiciclib in BT474 cancer cells. This was anomalous since the two blocking agents of HER2 and ER were expected to inhibit their crosstalk and achieve better responses. To explore the potential mechanisms, we investigated the crosstalk between HER2 and the ER. After degrading HER2 with pyrotinib, ER was found to relocate to the cell nucleus, enhancing the function of ER which was consistent with the findings of Kumar et al and Yang et al (*Kumar et al., 2002; Yang et al., 2004*). We believe that the anti-HER2 mediated ER redistribution caused the enhanced ER function, leading to the relatively low efficacy of the combination of pyrotinib and tamoxifen in the treatment of HER2^+^/HR^+^ cells. Moreover, we found that the introduction of dalpiciclib to pyrotinib significantly decreased the total and nuclear expression of ER, reversing the ER activation caused by pyrotinib (Figure2-figure supplement 1c). This may be the underlying mechanism by which the addition of a CDK4/6 inhibitor was more beneficial than the addition of pyrotinib and tamoxifen.

Furthermore, using RNA-seq and bioinformatics analyses, CALML5 was selected as a potential biomarker for responsiveness to anti-HER2 therapy combined with a CDK4/6 inhibitor. CALML5, known as calmodulin-like 5, is a skin-specific calcium-binding protein that is closely related to keratinocyte differentiation (*Mehul et al., 2001*) A previous study showed that the high expression of CALML5 was strongly associated with better survival in patients with head and neck squamous cell carcinomas (*Wirsing et al., 2021*). Misawa et al. (*Misawa et al., 2020*) reported that the methylation of CALML5, led to its downregulation, and this showed a correlation with HPV-associated oropharyngeal cancer. Moreover, the ubiquitination of CALML5 in the nucleus was found to play a role in the carcinogenesis of breast cancer in premenopausal women (*Debald et al., 2013*) Our results suggested that TPBC patients with positive CALML5 may benefit from the addition of CDK4/6 inhibitors in neoadjuvant therapy. However, the underlying mechanism of CALML5 in breast cancer requires further investigation.

In conclusion, our study showed the novel role of the CDK4/6 inhibitor in TPBC and provided evidence that CALML5 may be a potential biomarker in the prediction of the responsiveness of HER2^+^/HR^+^ breast cancer patients to CDK4/6 inhibitors.

## Materials and methods

### Clinical specimens

A total of 221 HR^+^/HER2^+^ patients who received adjuvant chemotherapy + anti HER2 therapy with or without endocrine therapy at the Shengjing Hospital of China Medical University were enrolled for the analysis of DFS and OS. A total of 198 HR^+^/HER2^+^ patients who received neoadjuvant therapy were enrolled in this study to evaluate the status of ER and CALML5, of which 26 patients were from the clinical trial (NCT04486911), 41 patients received anti-HER2 therapy plus chemotherapy, and 131 patients only received chemotherapy. The sample size was calculated based on the four interrelated statistics in the Null Hypothesis Significant Test (NHST): sample size, effect size, alpha level, and statistical efficacy. The clinical information and specimens were analyzed to determine the impact of endocrine therapy on prognosis.

The study was approved by the Institutional Ethics Committee and complied with the principles of the Declaration of Helsinki and Good Clinical Practice guidelines of the National Medical Products Administration of China. Informed consent was obtained from all the participants.

### Cell lines and cell cultures

BT474 and MCF7 were purchased from the American Type Culture Collection (ATCC, Manassas, VA, USA). The human triple-positive breast cancer cell line BT474 was cultured in RPMI1640 culture medium supplemented with 10% fetal bovine serum (FBS), and MCF7 cells were cultured in DMEM culture medium supplemented with 10% FBS.

### Chemicals and antibodies

Pyrotinib (SHR1258) and dalpiciclib (SHR6390) were kindly provided by Hengrui Medicine Co., Ltd. Tamoxifen (HY-13757A) was purchased from MCE company. Compounds were dissolved in dimethylsulfoxide (DMSO) at a concentration of 10 mM and stored at −20 °C for further use. The following antibodies were purchased from Cell Signaling Technology (Beverly, MA, USA): ER, p-HER2 (Tyr 1221/1222), HER2, p-Akt (Ser473), AKT, p-mTOR, mTOR, pRb (Ser 780), Rb, CDK4, CDK6, Lamin A, and GAPDH.

### Cell viability assays and drug combination studies

CCK cell viability assays were (Cofitt life science) used to quantify the inhibitory effect of the different treatments. Cells were seeded in 96-well plates at a density of 5000 cells/well and treated the next day with DMSO, pyrotinib, tamoxifen, dalpiciclib, or both drugs in combination for 48 h. The combination index (CI) values of different drugs were calculated using CompuSyn (ComboSyn Inc.). The CI values demonstrated synergistic (<1), additive (1–1.2), or antagonistic (>1.2) effects of the two-drug combinations. The drug sensitivity experiments were performed three times independently.

### Cell cycle analyses

The cells were starved in culture medium supplemented with 2% serum for 24 h before treatment. Treatments included DMSO (0.1%), pyrotinib (10 nM), dalpiciclib (8 μM), tamoxifen (5 μM), or different combinations of drugs. After treatment for 24 h, cells in different treating groups were trypsinized, washed with PBS, fixed in 70% ethanol, and incubated overnight at 4 °C. Next day, cells were collected, washed, and re-suspended in PBS at a concentration of 5 × 10^5^ cells/mL. The cell solutions were then incubated with a RNase and propidium iodide (PI) solution for 30 min at room temperature without exposure to light, and analyzed using a flow cytometer (BD FACS Calibur) according to the manufacturer’s instructions. This assay was performed in triplicates.

### Colony formation assays

Cells were seeded in 6-well plates at a density of 1000 cells/well. The cells were treated with DMSO (0.1%), pyrotinib (10 nM), tamoxifen (5 μM), dalpiciclib (8 μM), or a combination of the two or three agents. During the process, the culture medium was renewed every three days. After 14 days, the colonies were fixed and stained with crystal violet. Clusters of more than eight cells were counted as colonies. This assay was performed in triplicates independently.

### Transfection of the human HER2 plasmids

MCF7 cells were cultured in DMEM supplemented with 10% FBS. Human HER2 plasmids were purchased from Hanbio. The plasmids were mixed well with lipo3000 and p3000, according to the manufacturer’s instructions, and then added to the culture medium. Forty-eight hours after the transfection, the cells were fixed in formaldehyde and stained for the estrogen receptor.

### Western blot analysis

Cells and cancer tissues were lysed using a cell lysis buffer (Beyotime, Shanghai, China). The total proteins were extracted in a lysis buffer (Beyotime, Shanghai, China), and the nuclear proteins were extracted using a nuclear protein extraction kit (Beyotime), in which PMSF, protease, and phosphatase inhibitors were added. Protein concentrations were determined using a Pierce BCA Protein Assay Kit (Thermo Fisher Scientific, Waltham, MA, USA) according to the manufacturer’s instructions. The proteins from the cells and tissue lysates were separated using 10% SDS-PAGE and 6% SDS-PAGE, respectively, and then transferred to polyvinylidene fluoride (PVDF) membranes. The immunoreactive bands were detected using enhanced chemiluminescence (ECL). The western blot analysis was performed in triplicates independently.

### Immunofluorescence assays

The cellular localization of different proteins was detected using immunofluorescence. Briefly, the cells grown on glass coverslips were fixed in 4% paraformaldehyde at room temperature for 30 min. Cells were incubated with the respective primary antibodies for 1 h at room temperature, washed in PBS, and then incubated with 590-Alexa-(red) secondary antibodies (Molecular Probes, Eugene, OR, USA). We used 590-Alexa-phalloidin to localize the ER. The nuclei of the cells were stained with DAPI and color-coded in blue. The images were captured using an immunofluorescence microscope (Nikon Oplenic Lumicite 9000). The distribution ratio of ER was calculated manually by randomly chosen 5 views in 400magnification. The immunofluorescence assay was performed in triplicates independently.

### Immunohistochemical staining

The clinical samples were fixed in 4% formaldehyde, embedded in paraffin, and sectioned continuously at a thickness of 3 μm. The paraffin sections were deparaffinized with xylene and rehydrated using a graded ethanol series. They were then washed with tris-buffered saline (TBS). After these preparation procedures, the sections of each sample were incubated with the primary anti-ER antibody (Abcam Company, ab32063), anti-HER2 antibody (Abcam Company, ab134182), and anti-CALML5 antibody (Proteintech, 13059-1-AP) at 4 °C overnight. The next day, they were washed three times with TBS and incubated with a horseradish peroxidase (HRP)-conjugated secondary antibody (Gene Tech Co. Ltd., Shanghai, China) at 37 °C for 45 min, followed by immunohistochemical staining using a DAB kit (Gene Tech Co. Ltd.) for 5–10 min.

### Evaluation of the ER and HER2 statuses

The ER and HER2 statuses of patients who received neoadjuvant therapy were evaluated by a pathologist from a Shenjing affiliated hospital. The clinical specimens before and after the neoadjuvant therapy were evaluated. The analyses of the elevation or decline in ER statuses were based on these pathological reports.

### Gene enrichment analysis

Gene annotation data in the GO and KEGG databases and R language were used for the enrichment analysis. Only enrichment with q-values less than 0.05 were considered significant.

### GSEA

The hallmark gene sets in the Molecular Signatures Database were used for performing the GSEA; only gene sets with q-values less than 0.05 were considered significantly enriched.

### Statistical analysis

All the descriptive statistics (except the drug sensitivity assay in Figure 1a and b) were presented as the means ± standard deviations (SDs). The drug sensitivity assay in Figure 1a and b were presented as the means ± standard error of mean (SEM). The differences between the groups were analyzed by chi-squared or Student’s t tests. Kaplan-Meier methods were used to compute the survival analysis and *P*-value was obtained by log-rank test. The statistical analyses were performed using IBM SPSS version 22 (SPSS, Armonk, NY, USA) and GraphPad Prism version 7. The statistical significance of the differences between the test and control samples was assessed at significance thresholds of \**P* < 0.05, \*\**P* < 0.01, \*\*\**P* < 0.001, and \*\*\*\**P* < 0.0001.

## Authors Contributions

J.B., Y.Z., L.S., X.Q., Y.W., X.J., D.W., H.L., and Q.M. conceptualized the study, performed the experiments, and analyzed the data. B.K. performed the bioinformatic analysis. Y.Z. and N.N. provided the clinical data and samples. C.L. designed the entire study and wrote the manuscript.

## Conflict of interest

The authors declare no conflicts of interests. H.L. is the employee of Jiangsu Hengrui Pharmaceuticals Co., Ltd, no other potential conflicts of interest were reported. The other authors have no competing interests to declare.

## Funding

This study was supported by the National Natural Science Foundation of China (#U20A20381, #81872159)

## Supplementary

**Figure 1-figure supplement 1.**
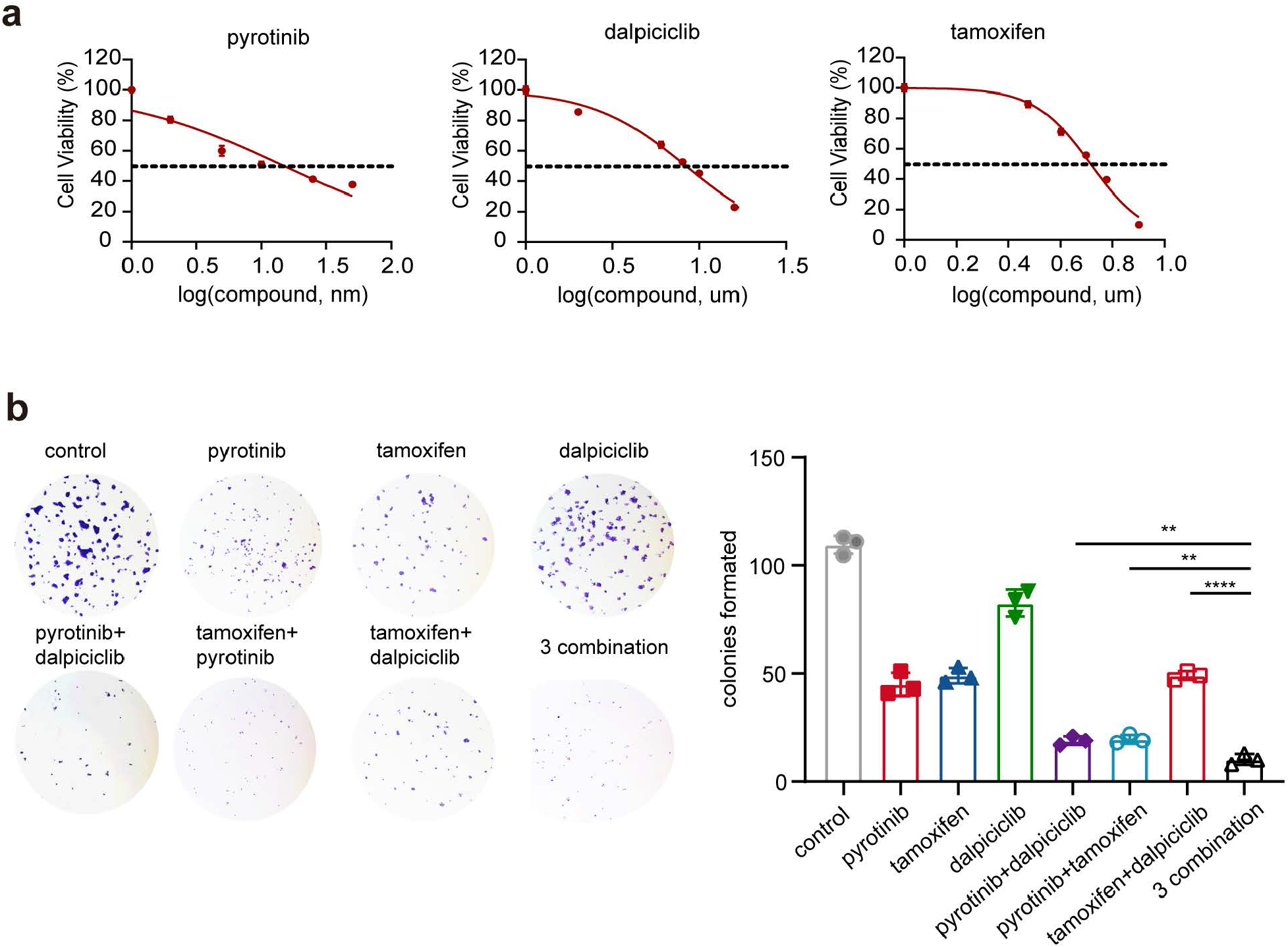
a: Drug sensitivity analysis of pyrotinib, tamoxifen and dalpiciclib in BT474 cells. (Data was presented as mean ± SDs, all the assays were performed independently in triplicates). b-c: Colony formation assay of BT474 cells treated with different drugs. (Data was presented as mean ± SDs, \*\**P*<0.01 and \*\*\*\**P*<0.0001 using Student’s *t*-test; all the assays were performed independently in triplicates).

**Figure 2-figure supplement 1.**
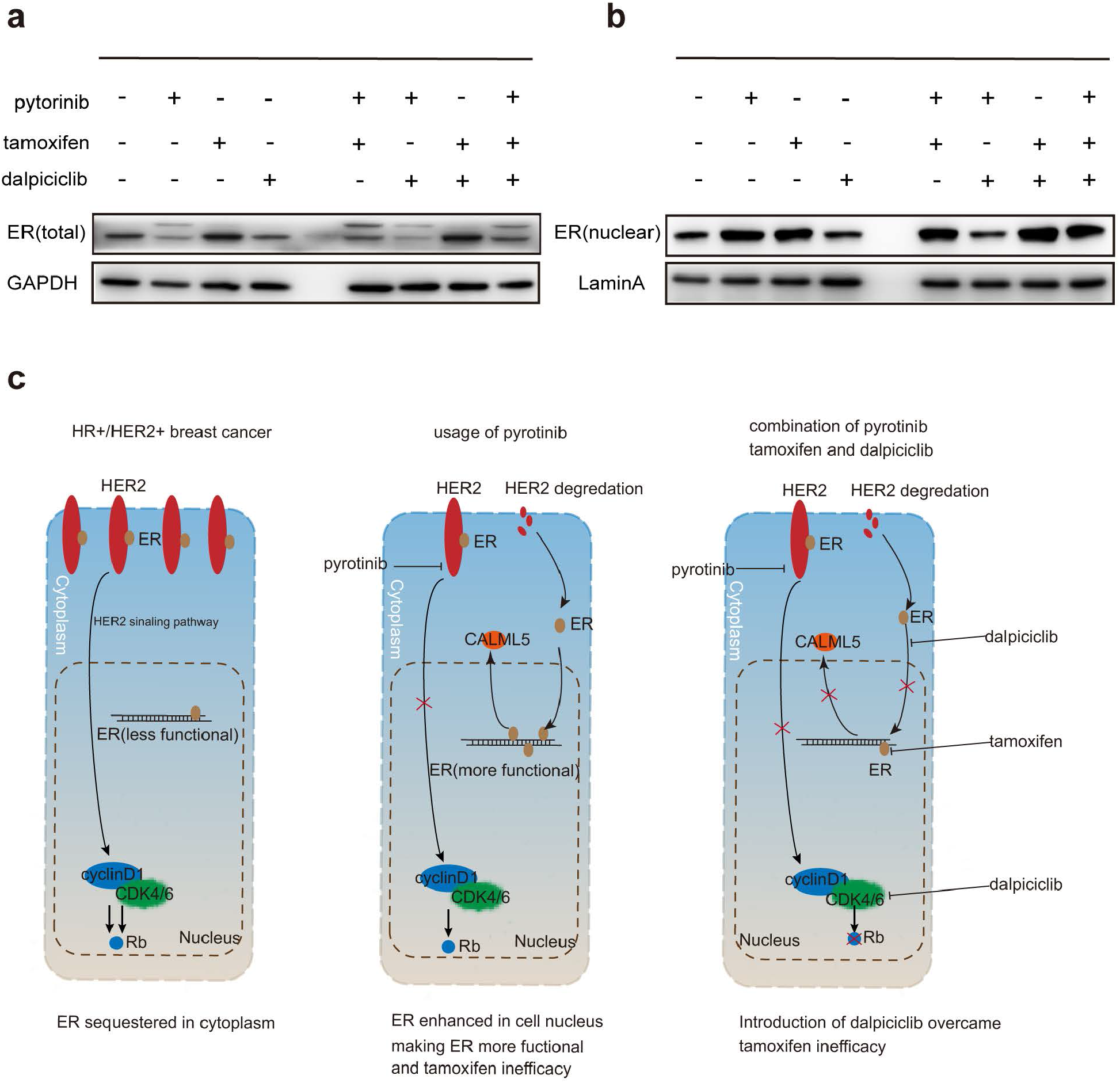
a-b: Total ER expression and nuclear ER expression in BT474 cells treated with different drugs. (This assay was performed in triplicates independently). c: The introduction of dalpiciclib to pyrotinib could significantly decrease the total and nuclear expression of ER, thus reversed the ER activation caused by pyrotinib and CALML5 could be served as a potential marker of ER activation after the treatment of pyrotinib.

## Source data

**Figure 2 source data 1**

The clinical pathological data of patients who received neoadjuvant chemotherapy + anti HER2 therapy

**Figure 2 source data 2**

The clinical pathological data of patients who received neoadjuvant chemotherapy only.

**Figure 3 source data 1**

RNA-sequencing data of BT474 cells treated with different drug combination.

**Figure 4 source data 1**

Original files of western blot analysis in Figure 4 a.

**Figure 2-figure supplement 1 source data 1**

Original files of western blot analysis in Figure 2-figure supplement 1 a-b.

## Notes

### Competing Interest Statement

The authors have declared no competing interest.

## References

Brandao, M, Caparica, R, Malorni, L, Prat, A, Carey, LA, and Piccart, M. 2020. What Is the Real Impact of Estrogen Receptor Status on the Prognosis and Treatment of HER2-Positive Early Breast Cancer? Clin Cancer Res 26, 2783–2788. DOI: https://doi.org/10.1158/1078-0432.CCR-19-2612, PMID:32046997.

Cameron, D, Piccart-Gebhart, MJ, Gelber, RD, Procter, M, Goldhirsch, A, de Azambuja, E, Castro, G, Jr., Untch, M, Smith, I, Gianni, L, Baselga, J, Al-Sakaff, N, Lauer, S, McFadden, E, Leyland-Jones, B, Bell, R, Dowsett, M, Jackisch, C, and Herceptin Adjuvant Trial Study, T. 2017. 11 years’ follow-up of trastuzumab after adjuvant chemotherapy in HER2-positive early breast cancer: final analysis of the HERceptin Adjuvant (HERA) trial. Lancet 389, 1195–1205. DOI: https://doi.org/10.1016/S0140-6736(16)32616-2, PMID:28215665.

Carey, LA, Berry, DA, Cirrincione, CT, Barry, WT, Pitcher, BN, Harris, LN, Ollila, DW, Krop, IE, Henry, NL, Weckstein, DJ, Anders, CK, Singh, B, Hoadley, KA, Iglesia, M, Cheang, MC, Perou, CM, Winer, EP, and Hudis, CA. 2016. Molecular Heterogeneity and Response to Neoadjuvant Human Epidermal Growth Factor Receptor 2 Targeting in CALGB 40601, a Randomized Phase III Trial of Paclitaxel Plus Trastuzumab With or Without Lapatinib. J Clin Oncol 34, 542–549. DOI: https://doi.org/10.1200/JCO.2015.62.1268, PMID:26527775.

Chou, TC, and Talalay, P. 1984. Quantitative analysis of dose-effect relationships: the combined effects of multiple drugs or enzyme inhibitors. Adv Enzyme Regul 22, 27–55. DOI: https://doi.org/10.1016/0065-2571(84)90007-4, PMID:6382953.

Cortazar, P, Zhang, L, Untch, M, Mehta, K, Costantino, JP, Wolmark, N, Bonnefoi, H, Cameron, D, Gianni, L, Valagussa, P, Swain, SM, Prowell, T, Loibl, S, Wickerham, DL, Bogaerts, J, Baselga, J, Perou, C, Blumenthal, G, Blohmer, J, Mamounas, EP, et al. 2014. Pathological complete response and long-term clinical benefit in breast cancer: the CTNeoBC pooled analysis. Lancet 384, 164–172. DOI: https://doi.org/10.1016/S0140-6736(13)62422-8, PMID:24529560.

Debald, M, Schildberg, FA, Linke, A, Walgenbach, K, Kuhn, W, Hartmann, G, and Walgenbach-Brunagel, G. 2013. Specific expression of k63-linked ubiquitination of calmodulin-like protein 5 in breast cancer of premenopausal patients. J Cancer Res Clin Oncol 139, 2125–2132. DOI: https://doi.org/10.1007/s00432-013-1541-y, PMID:24146193.

Gianni, L, Bisagni, G, Colleoni, M, Del Mastro, L, Zamagni, C, Mansutti, M, Zambetti, M, Frassoldati, A, De Fato, R, Valagussa, P, and Viale, G. 2018. Neoadjuvant treatment with trastuzumab and pertuzumab plus palbociclib and fulvestrant in HER2-positive, ER-positive breast cancer (NA-PHER2): an exploratory, open-label, phase 2 study. Lancet Oncol 19, 249–256. DOI: https://doi.org/10.1016/S1470-2045(18)30001-9, PMID:29326029.

Gianni, L, Pienkowski, T, Im, YH, Roman, L, Tseng, LM, Liu, MC, Lluch, A, Staroslawska, E, de la Haba-Rodriguez, J, Im, SA, Pedrini, JL, Poirier, B, Morandi, P, Semiglazov, V, Srimuninnimit, V, Bianchi, G, Szado, T, Ratnayake, J, Ross, G, and Valagussa, P. 2012. Efficacy and safety of neoadjuvant pertuzumab and trastuzumab in women with locally advanced, inflammatory, or early HER2-positive breast cancer (NeoSphere): a randomised multicentre, open-label, phase 2 trial. Lancet Oncol 13, 25–32. DOI: https://doi.org/10.1016/S1470-2045(11)70336-9, PMID:22153890.

Goel, S, Wang, Q, Watt, AC, Tolaney, SM, Dillon, DA, Li, W, Ramm, S, Palmer, AC, Yuzugullu, H, Varadan, V, Tuck, D, Harris, LN, Wong, KK, Liu, XS, Sicinski, P, Winer, EP, Krop, IE, and Zhao, JJ. 2016. Overcoming Therapeutic Resistance in HER2-Positive Breast Cancers with CDK4/6 Inhibitors. Cancer Cell 29, 255–269. DOI: https://doi.org/10.1016/j.ccell.2016.02.006, PMID:26977878.

Harbeck, N, Gluz, O, Christgen, M, Kates, RE, Braun, M, Kuemmel, S, Schumacher, C, Potenberg, J, Kraemer, S, Kleine-Tebbe, A, Augustin, D, Aktas, B, Forstbauer, H, Tio, J, von Schumann, R, Liedtke, C, Grischke, EM, Schumacher, J, Wuerstlein, R, Kreipe, HH, et al. 2017. De-Escalation Strategies in Human Epidermal Growth Factor Receptor 2 (HER2)-Positive Early Breast Cancer (BC): Final Analysis of the West German Study Group Adjuvant Dynamic Marker-Adjusted Personalized Therapy Trial Optimizing Risk Assessment and Therapy Response Prediction in Early BC HER2- and Hormone Receptor-Positive Phase II Randomized Trial-Efficacy, Safety, and Predictive Markers for 12 Weeks of Neoadjuvant Trastuzumab Emtansine With or Without Endocrine Therapy (ET) Versus Trastuzumab Plus ET. J Clin Oncol 35, 3046–3054. DOI: https://doi.org/10.1200/JCO.2016.71.9815, PMID:28682681.

Kumar, R, Wang, RA, Mazumdar, A, Talukder, AH, Mandal, M, Yang, Z, Bagheri-Yarmand, R, Sahin, A, Hortobagyi, G, Adam, L, Barnes, CJ, and Vadlamudi, RK. 2002. A naturally occurring MTA1 variant sequesters oestrogen receptor-alpha in the cytoplasm. Nature 418, 654–657. DOI: https://doi.org/10.1038/nature00889, PMID:12167865.

Loi, S, Dafni, U, Karlis, D, Polydoropoulou, V, Young, BM, Willis, S, Long, B, de Azambuja, E, Sotiriou, C, Viale, G, Ruschoff, J, Piccart, MJ, Dowsett, M, Michiels, S, and Leyland-Jones, B. 2016. Effects of Estrogen Receptor and Human Epidermal Growth Factor Receptor-2 Levels on the Efficacy of Trastuzumab: A Secondary Analysis of the HERA Trial. JAMA Oncol 2, 1040–1047. DOI: https://doi.org/10.1001/jamaoncol.2016.0339, PMID:27100299.

Maguire, R, McCann, L, Kotronoulas, G, Kearney, N, Ream, E, Armes, J, Patiraki, E, Furlong, E, Fox, P, Gaiger, A, McCrone, P, Berg, G, Miaskowkski, C, Cardone, A, Orr, D, Flowerday, A, Katsaragakis, S, Darley, A, Lubowitzki, S, Harris, J, et al. 2021. Real time remote symptom monitoring during chemotherapy for cancer: European multicentre randomised controlled trial (eSMART). BMJ 374, n1647. DOI: https://doi.org/10.1136/bmj.n1647, PMID:34289996.

Mehul, B, Bernard, D, and Schmidt, R. 2001. Calmodulin-like skin protein: a new marker of keratinocyte differentiation. J Invest Dermatol 116, 905–909. DOI: https://doi.org/10.1046/j.0022-202x.2001.01376.x, PMID:11407979.

Misawa, K, Imai, A, Matsui, H, Kanai, A, Misawa, Y, Mochizuki, D, Mima, M, Yamada, S, Kurokawa, T, Nakagawa, T, and Mineta, H. 2020. Identification of novel methylation markers in HPV-associated oropharyngeal cancer: genome-wide discovery, tissue verification and validation testing in ctDNA. Oncogene 39, 4741–4755. DOI: https://doi.org/10.1038/s41388-020-1327-z, PMID:32415241.

Moja, L, Tagliabue, L, Balduzzi, S, Parmelli, E, Pistotti, V, Guarneri, V, and D’Amico, R. 2012. Trastuzumab containing regimens for early breast cancer. Cochrane Database Syst Rev, CD006243. DOI: https://doi.org/10.1002/14651858.CD006243.pub2, PMID:22513938.

Pascual, J, Lim, JSJ, Macpherson, IR, Armstrong, AC, Ring, A, Okines, AFC, Cutts, RJ, Herrera-Abreu, MT, Garcia-Murillas, I, Pearson, A, Hrebien, S, Gevensleben, H, Proszek, PZ, Hubank, M, Hills, M, King, J, Parmar, M, Prout, T, Finneran, L, Malia, J, et al. 2021. Triplet Therapy with Palbociclib, Taselisib, and Fulvestrant in PIK3CA-Mutant Breast Cancer and Doublet Palbociclib and Taselisib in Pathway-Mutant Solid Cancers. Cancer Discov 11, 92–107. DOI: https://doi.org/10.1158/2159-8290.CD-20-0553, PMID:32958578.

Perou, CM, Sorlie, T, Eisen, MB, van de Rijn, M, Jeffrey, SS, Rees, CA, Pollack, JR, Ross, DT, Johnsen, H, Akslen, LA, Fluge, O, Pergamenschikov, A, Williams, C, Zhu, SX, Lonning, PE, Borresen-Dale, AL, Brown, PO, and Botstein, D. 2000. Molecular portraits of human breast tumours. Nature 406, 747–752. DOI: https://doi.org/10.1038/35021093, PMID:10963602.

Rimawi, M, Cecchini, R, Rastogi, P, Geyer, C, Fehrenbacher, L, Stella, P, Dayao, Z, Rabinovitch, R, Dyar, S, Flynn, P, Baez-Diaz, L, Paik, S, Swain, S, Mamounas, E, Osborne, C, and Wolmark, N. 2017. Abstract S3-06: A phase III trial evaluating pCR in patients with HR+, HER2-positive breast cancer treated with neoadjuvant docetaxel, carboplatin, trastuzumab, and pertuzumab (TCHP) +/-estrogen deprivation: NRG Oncology/NSABP B-52. Cancer Research 77, S3-06-S03-06. DOI: https://doi.org/10.1158/1538-7445.sabcs16-s3-06.

Saura, C, Garcia-Saenz, JA, Xu, B, Harb, W, Moroose, R, Pluard, T, Cortes, J, Kiger, C, Germa, C, Wang, K, Martin, M, Baselga, J, and Kim, SB. 2014. Safety and efficacy of neratinib in combination with capecitabine in patients with metastatic human epidermal growth factor receptor 2-positive breast cancer. J Clin Oncol 32, 3626–3633. DOI: https://doi.org/10.1200/JCO.2014.56.3809, PMID:25287822.

Schneeweiss, A, Chia, S, Hickish, T, Harvey, V, Eniu, A, Hegg, R, Tausch, C, Seo, JH, Tsai, YF, Ratnayake, J, McNally, V, Ross, G, and Cortes, J. 2013. Pertuzumab plus trastuzumab in combination with standard neoadjuvant anthracycline-containing and anthracycline-free chemotherapy regimens in patients with HER2-positive early breast cancer: a randomized phase II cardiac safety study (TRYPHAENA). Ann Oncol 24, 2278–2284. DOI: https://doi.org/10.1093/annonc/mdt182, PMID:23704196.

Slamon, DJ, Clark, GM, Wong, SG, Levin, WJ, Ullrich, A, and McGuire, WL. 1987. Human breast cancer: correlation of relapse and survival with amplification of the HER-2/neu oncogene. Science 235, 177–182. DOI: https://doi.org/10.1126/science.3798106, PMID:3798106.

Tzahar, E, Waterman, H, Chen, X, Levkowitz, G, Karunagaran, D, Lavi, S, Ratzkin, BJ, and Yarden, Y. 1996. A hierarchical network of interreceptor interactions determines signal transduction by Neu differentiation factor/neuregulin and epidermal growth factor. Mol Cell Biol 16, 5276–5287. DOI: https://doi.org/10.1128/MCB.16.10.5276, PMID:8816440.

Wang, YC, Morrison, G, Gillihan, R, Guo, J, Ward, RM, Fu, X, Botero, MF, Healy, NA, Hilsenbeck, SG, Phillips, GL, Chamness, GC, Rimawi, MF, Osborne, CK, and Schiff, R. 2011. Different mechanisms for resistance to trastuzumab versus lapatinib in HER2-positive breast cancers--role of estrogen receptor and HER2 reactivation. Breast Cancer Res 13, R121. DOI: https://doi.org/10.1186/bcr3067, PMID:22123186.

Wirsing, AM, Bjerkli, IH, Steigen, SE, Rikardsen, O, Magnussen, SN, Hegge, B, Seppola, M, Uhlin-Hansen, L, and Hadler-Olsen, E. 2021. Validation of Selected Head and Neck Cancer Prognostic Markers from the Pathology Atlas in an Oral Tongue Cancer Cohort. Cancers (Basel) 13. DOI: https://doi.org/10.3390/cancers13102387, PMID:34069237.

Yang, Z, Barnes, CJ, and Kumar, R. 2004. Human epidermal growth factor receptor 2 status modulates subcellular localization of and interaction with estrogen receptor alpha in breast cancer cells. Clin Cancer Res 10, 3621–3628. DOI: https://doi.org/10.1158/1078-0432.CCR-0740-3, PMID:15173068.

Zhang, K, Hong, R, Kaping, L, Xu, F, Xia, W, Qin, G, Zheng, Q, Lu, Q, Zhai, Q, Shi, Y, Yuan, Z, Deng, W, Chen, M, and Wang, S. 2019. CDK4/6 inhibitor palbociclib enhances the effect of pyrotinib in HER2-positive breast cancer. Cancer Lett 447, 130–140. DOI: https://doi.org/10.1016/j.canlet.2019.01.005, PMID:30677445.

